# Viromes of Antarctic fish resembles the diversity found at lower latitudes

**DOI:** 10.1101/2024.04.29.591789

**Authors:** Rebecca M. Grimwood, Stephanie J. Waller, Janelle R. Wierenga, Lauren Lim, Jeremy Dubrulle, Edward C. Holmes, Jemma L. Geoghegan

**Affiliations:** Department of Microbiology and Immunology, University of Otago, Dunedin, 9016, New Zealand; Sydney Institute for Infectious Diseases, School of Medical Sciences, The University of Sydney, NSW, 2006, Australia; Institute of Environmental Science and Research, Wellington, 5018, New Zealand

**Keywords:** Antarctica, Ross Sea, fish, viromes, evolution, genomes

## Abstract

Antarctica harbours some of the most isolated and extreme environments on Earth, concealing a largely unexplored and unique component of the global animal virosphere. To understand the diversity and evolutionary histories of viruses in these polar species we determined the viromes of 11 Antarctic fish species with 248 samples collected from the Ross Sea region spanning the Perciformes, Gadiformes, and Scorpaeniformes orders. The continent’s shift southward and cooling temperatures over 20 million years ago led to a reduction in biodiversity and subsequent radiation of some marine fauna, such as the notothenioid fishes. Despite decreased host species richness in polar regions, we revealed a surprisingly complex virome diversity in Ross Sea fish, with the types and numbers of viruses per host species and individuals sampled comparable to that of fish in warmer marine environments with higher host community diversity. We also observed a higher number of closely related viruses likely representing instances of recent and historic host-switching events among Perciformes (all notothenioids) than in the Gadiformes, suggesting that rapid speciation events within this order generated closely related host species with few genetic barriers to cross-species transmission. Additionally, we identified novel genomic variation in an arenavirus with a split nucleoprotein sequence containing a stable helical structure, indicating potential adaptation of viral proteins to extreme temperatures. These findings enhance our understanding of virus evolution and virus-host interactions in response to environmental shifts, especially in less diverse ecosystems more vulnerable to the impacts of anthropogenic and climate changes.

## Introduction

Antarctica’s icy waters hide some of the world’s most isolated and hostile environments, and with it, its aquatic life shields a unique and largely unexplored pocket of the vertebrate virosphere. As the continent shifted southward over 20 million years ago, the cooling of the Southern Ocean towards an increasingly polar climate saw a reduction in the temperate Eocene fauna that previously inhabited Antarctica and opened increasingly extreme and isolated niches, reshaping the diversity of its marine life (Daane & Detrich, 2022; Eastman, 2005). Water temperatures in the Southern Ocean consistently sit below the freezing point of fish blood (Littlepage, 2013). Ray-finned fishes from the Notothenioidei, *Liparidae*, and Zoarcoidei account for most of the fish diversity in Antarctica (Eastman, 2005), and unique adaptations – such as antifreeze glycoproteins (AFGPs) in their blood – allowed them to prevail in these sub-zero climates where other temperate species could not (Devries, 1971). Subsequent specialisation to the changing environment, including instances of radiation, particularly of the notothenioid species (order: Perciformes), has created a taxonomically restricted modern ichthyofauna (Clarke & Johnston, 1996; Eastman & Clarke, 1998). Today, there are approximately 320 known species of Antarctic fish (Eastman, 2005), a relatively small number given that fish compose more than 50% of Earth’s vertebrate species (Nelson, 2006). Notothenioids alone have radiated to over 130 species and make up over 90% of the fish biomass in the Southern Ocean (Daane & Detrich, 2022; Eastman & Hubold, 1999). Radiation of Antarctic fishes has also resulted in the highest rates of endemism of any isolated marine environment (Eastman & Clarke, 1998). Such species not only make for fascinating models of evolution (Seehausen, 2006; Thacker et al., 2021), but also for studying their associated microbial flora, particularly viruses (Costa, Ronco, et al., 2023; Perry et al., 2022). Still, much of the focus on the viruses of Antarctic species to date has revolved around birds, seals (Smeele et al., 2018; Varsani et al., 2017), environmental samples (Aguirre de Cárcer et al., 2016; Gong et al., 2018), as well as Trematomus fish (Kraberger et al., 2022)(Kraberger et al., 2022)(Kraberger et al., 2022).

Growing attention to these polar species has also left them vulnerable to several threats. Antarctic fish, and their AFGPs, are scientifically, economically, and commercially important (Collins et al., 2010; Eskandari et al., 2020). However, overfishing in the last 50 years has induced concerns over population recovery, bycatch, and competition from marine predators. This is particularly acute for highly sought-after species like the Patagonian toothfish (*Dissostichus eleginoides*) and common bycatch such as grenadier and *Antimora rostrata* (Collins et al., 2010; Smith et al., 2012) for which little is known about their biology and ecology. Antarctic fish are also sensitive to environmental change. The Southern Ocean has been warming considerably faster than the global ocean (Gille, 2002) and unlike Antarctica’s specious tropical counterparts, the lack of biodiversity in this region leaves it without a protective buffer making it less resilient to climatic and anthropogenic changes (Duffy et al., 2016). Mortality of fish due to viral infections and modulation of host immune responses has also been shown to be affected by increasing temperatures (Jung et al., 2017; Páez et al., 2021). Hence, a better understanding of the biology, ecology, and microbes of polar fish may be important in the management and protection of these species in the face of these growing issues.

Species richness increases towards the equator (Rabosky et al., 2018) and this diversity has recently been shown to be equally manifested in that of fish viromes in these warmer waters (Costa, Bellwood, et al., 2023). Due to the relatively restricted and specialist aquatic taxonomy in the Southern Ocean, it may be expected that the viromes of these species are similarly restricted or divergent in comparison to those in more equatorial ecosystems. The Ross Sea is the largest embayment in Antarctica at 180° S and contains the world’s largest marine protected area (Cummings et al., 2021; Smith et al., 2012). At least 80 fish species reside in the Ross Sea, dominated by the notothenioids (making up more than 60% of species) and a number of non-notothenioid species (Eastman & Hubold, 1999). To better understand the virome of these lesser-studied aquatic species we sampled Antarctic fishes from the Ross Sea, spanning the Perciformes, Gadiformes, and Scorpaeniformes orders. Our aim was to use metatranscriptomic sequencing to examine their virome diversity and determine how historic oceanographic, evolutionary, and climatic changes may have shaped the diversity and evolution of these viruses compared to those of more temperature or tropical species.

## Methods

### Ethics

Samples were collected under the San Aotea II 2022 AMLR permit, the Ross Sea data collection plan, the Ross Sea shelf survey plan, and following the relevant Commission for the Conservation of Antarctic Marine Living Resources (CCAMLR) Conservation Measures.

### Gill sample collection

Antarctic fish were collected between early December 2021 and mid January 2022 by the San Aotea II during commercial fishing in the Ross Sea and research fishing on the Ross Sea shelf survey at depths of 522 to 1810 m. Species were identified where possible or placed into the next most descriptive taxonomic group (family or genus). Importantly, all of the fish were in good condition with no obvious signs of disease. Whole fish heads were frozen and stored at -4°C ready for dissection of gill tissue.

Frozen heads were partially defrosted and approximately 0.5 – 1 cm^3^ of gill tissue was dissected using sterilised scalpels and forceps and stored in 1 mL of RNA*later* (ThermoFisher Scientific) before storage at -80°C. Gill samples were taken from between four and 23 individuals per species group for a total of 248 samples from three orders: Perciformes (n=118) (*Chionobathyscus dewitti*, *Trematomus loennbergii*, *Trematomus lepidorhinus*, *Pogonophryne barsukovi*, *Pogonophryne immaculata*, and *Pogonophryne scotti*), Gadiformes (n=121) (*Macrourus caml*, *Macrourus whitsoni*, *Antimora rostrata*, and *Muraenolepis spp.*), and Scorpaeniformes (n=9) (*Zoarcidae spp.*) (see Table 1).

**Table 1.** Overview of sampling libraries.

| Library | Host species name/group | Taxonomic order | Number of fish | Body mass (kg) |
| --- | --- | --- | --- | --- |
| MRL_L | *Muraenolepis spp.* | Gadiformes | 11 | 0.51 – 0.74 |
| MRL_H | *Muraenolepis spp.* | Gadiformes | 8 | 0.77 – 1.01 |
| ANT_L | *Antimora rostrata* | Gadiformes | 12 | 0.98 – 1.91 |
| ANT_H | *Antimora rostrata* | Gadiformes | 11 | 2.02 – 3.72 |
| WGR_L | *Macrourus whitsoni* | Gadiformes | 18 | 0.21 – 0.58 |
| WGR_H | *Macrourus whitsoni* | Gadiformes | 20 | 0.6 – 1.28 |
| QMC_L | *Macrourus caml* | Gadiformes | 23 | 0.67 – 1.26 |
| QMC_H | *Macrourus caml* | Gadiformes | 18 | 1.24 – 2.25 |
| LVD_L | *Zoarcidae* | Scorpaeniformes | 5 | 0.04 – 0.1 |
| LVD_H | *Zoarcidae* | Scorpaeniformes | 4 | 0.11 – 0.22 |
| TRD_L | *Trematomus lepidorhinus* | Perciformes | 11 | 0.14 – 0.32 |
| TRD_H | *Trematomus lepidorhinus* | Perciformes | 8 | 0.33 – 0.5 |
| TLO_L | *Trematomus loennbergii* | Perciformes | 10 | 0.19 – 0.35 |
| TLO_H | *Trematomus loennbergii* | Perciformes | 7 | 0.36 – 0.52 |
| CHW_L | *Chionobathyscus dewitti* | Perciformes | 21 | 0.17 – 0.29 |
| CHW_H | *Chionobathyscus dewitti* | Perciformes | 19 | 0.3 – 0.68 |
| SZT_L | *Pogonophryne scotti* | Perciformes | 5 | 0.19 – 0.26 |
| SZT_H | *Pogonophryne scotti* | Perciformes | 5 | 0.3 – 0.43 |
| PIM_L | *Pogonophryne immaculata* | Perciformes | 9 | 0.15 – 0.24 |
| PIM_H | *Pogonophryne immaculata* | Perciformes | 9 | 0.26 – 0.36 |
| BVK_L | *Pogonophryne barsukovi* | Perciformes | 7 | 0.26 – 0.38 |
| BVK_H | *Pogonophryne barsukovi* | Perciformes | 7 | 0.4 – 0.5 |

### RNA extraction and sequencing

For RNA extractions, the RNeasy Plus Mini Kit (Qiagen) protocol was followed for all samples. Frozen gill tissue was initially prepared by partially thawing and ∼10 mg was taken using sterilised scalpels and forceps and placed into RLT lysis buffer containing 1% β-mercaptoethanol and 0.5% (v/v) Reagent DX. A Qiagen TissueRupture was used to homogenise the tissue in the lysis buffer for around 30 seconds per sample. The resulting homogenates were centrifuged to remove residual tissue and the protocol was resumed to extract whole RNA. RNA concentrations were quantified using a NanoDrop Spectrophotometer (ThermoFisher).

RNA (5-20 μL) from each individual was pooled by species and into either “low” (L) or “high” (H) body mass (kg) for a total of 22 library pools containing RNA from four to 23 fish each (see Table 1).

Illumina Stranded Total RNA Prep with Ribo-Zero plus (Illumina) was used for library preparation of the 22 pooled RNA libraries and these were subjected to total RNA paired-end sequencing on the Illumina NovaSeq 6000 platform, generating 150 bp reads.

### Fish gill metatranscriptome assembly and annotation

Trinity RNA-seq (v2.11) (Grabherr et al., 2011) was used for *de novo* assembly of the gill meta-transcriptomes. Default parameters for paired-end read inputs were used with the additional “trimmomatic’ flag option to perform pre-assembly quality trimming of the reads. Trinity contigs from each library were annotated at both the nucleotide level against NCBI’s nucleotide (nt) database using the BLASTn algorithm (Camacho et al., 2009) and at the amino acid level against the non-redundant (nr) protein database using DIAMOND BLASTx (v2.02.2) (Buchfink et al., 2021).

### Vertebrate virus discovery and abundance estimation

Contigs annotated as being potentially viral in origin in one or both of the nucleotide or protein search outputs with e-values of less than 1×10^-10^ were checked with additional blastn and blastp searches using the online BLAST server (https://blast.ncbi.nlm.nih.gov/Blast.cgi) to eliminate false positive hits. Sequences with e-values greater than 1×10^-10^ or where the top blastn or blastp hits included non-viral or non-vertebrate viral species based on their taxonomic assignments and/or proposed hosts (e.g. bacteriophages, environmental metagenome-, fungi-, plant-, or invertebrate-associated viruses) were excluded from further analyses. Putative vertebrate virus sequences were considered if their top blastp hits were to those of other previously known vertebrate viruses. DNA virus sequences were also checked with additional blastn searches to eliminate those sequences with possible homology to fish genomic DNA and exclude possible endogenous viral elements (EVEs).

Abundance estimations of contigs were generated using the “align and estimate” module within Trinity with the “prep reference” flag set, RNA-seq by Expectation-Maximization (RSEM) (B. Li & Dewey, 2011) as the abundance estimation method, and Bowtie 2 (Langmead & Salzberg, 2012) as the alignment method. Contig abundances were standardised for inter-library comparisons by dividing RSEM abundance counts by the total sequencing read depth in their respective libraries. Standardised abundances were converted to reads-per-million (RPM) by multiplying by one million. To reduce incorrect assignment of viruses to libraries due to index-hopping, virus sequences sharing more than 99% nucleotide identity with sequences in other libraries occurring at abundances less than 0.1% of the highest abundance for that virus across all libraries were considered contamination and were removed from subsequent analyses.

For comparison of the presence and abundance of viruses between host taxonomic groups, abundances from each of the two body mass libraries per host group were merged. Abundances of individual viruses were compared alongside the standardised abundance of the stably expressed host gene, 40S Ribosomal Protein S13 (RPS13). RPS13 shows minimal inter- and intra-tissue expression level variation, making it a suitable reference host gene for transcript abundance comparisons (Robichaux et al., 2016). Where full genomes or genome segments of novel viruses were suspected due to the presence of full open reading frames (ORFs) of the expected lengths for viruses in their respective families, all novel virus segments for that virus were aligned with those from their closest known relatives for confirmation.

### Phylogenetic analysis

Inferences about the evolutionary relationships of the viruses documented here were made using maximum likelihood trees generated with IQ-TREE (v1.6.12) (Nguyen et al., 2015). We assumed that the viruses identified here that clustered with known viruses from other fish or vertebrate host species in their respective phylogenies were likely to be replicating in the fish host directly, rather than being associated with fish diet, microbiome or environment. All phylogenies were generated using amino acid sequences containing the highly conserved polymerase genes (either the RNA-dependent RNA polymerase (RdRp) for RNA viruses or the DNA-dependent DNA polymerase (DdDp) of DNA viruses). Viruses that could be assigned down to at least order level based on shared sequence identity with their closest relatives identified by BLAST searches were collated with a range of viruses from their respective taxonomies retrieved from NCBI Taxonomy (https://www.ncbi.nlm.nih.gov/Taxonomy/Browser/wwwtax.cgi) and aligned using the MAFFT (v7.450) (Katoh & Standley, 2013) L-INS-i algorithm. The exception were viruses from the *Nackednaviridae* and *Hepadnaviridae* families, which were aligned together with sequences from (Lauber et al., 2017) rather than from NCBI Taxonomy. Alignments were manually inspected in Geneious Prime (v2023.2.1) (https://www.geneious.com/) to observe the expected alignment of conserved amino acid motifs and ambiguously aligned regions were trimmed using trimAl (V1.2) (Capella-Gutiérrez et al., 2009) with the “automated1” flag set, or in the case of the *Picornaviridae*, with the flags “gt” (gap threshold) set to 0.9 and “cons” (variable conserve value) set to 50 in order to prevent excessive trimming. Sequence alignments for each taxonomic group were used to estimate phylogenies using IQ-TREE with the LG amino acid substitution model and 1000 ultra-fast bootstrapping replicates (Hoang et al., 2018). The “alrt” flag was also added to perform 1000 bootstrap replicates for the SH-like approximate likelihood ratio test (Guindon et al., 2010).

Phylogenetic trees were annotated in FigTree (v1.4.4) (http://tree.bio.ed.ac.uk/software/figtree/) and rooted at their midpoints.

For viruses of the *Arenaviridae* found in pools of *Trematomous* fish in this study and through data mining (see below), phylogenetic trees were also produced using amino acid sequences of nucleoproteins using the same method above.

### Data mining for arenaviruses and virus genome segment recovery

To identify further arenaviruses that may have unique genomic and structural variations present in the NCBI Transcriptome Shotgun Assembly (TSA) database, full-length amino acid sequences of the L protein (polymerase), segmented nucleoprotein, and glycoprotein from Trematomous arenavirus (PP590693 and PP590768-9) were used as bait to screen all vertebrate transcriptome assemblies

(taxonomic identifier: 7742), excluding *Homo sapiens* (taxonomic identifier: 9606), using the online translated Basic Local Alignment nucleotide (tblastn) search tool. The BLOSUM45 matrix was used to increase the chance of finding highly divergent viruses. Any putative virus hits to the “bait” sequences were queried using further blastp searches for confirmation. Raw read data associated with all positive hits were recovered from the Short Read Archive (SRA) as follows: SRR3184758 (resting phase testis of *Channa punctata*), SRR12526228 (RNAseq of *Coregonus artedi*), and SRR2912518 (transcriptome sequencing of white sucker (*Catostomus commersonii*)). Trinity RNA-seq (see Fish gill metatranscriptome assembly and annotation) for SRR318478 and MEGAHIT (v1.2.9) (D. Li et al., 2015) for SRR12526228 and SRR2912518 with default parameters were used to reassemble transcriptomes of the mined host data in an attempt to recover all possible genome segments for each new arenavirus identified. Assembled transcriptomes were screened against the nr protein database using DIAMOND BLASTx searches and the outputs were manually screened for the sequences of interest.

### Nucleoprotein structural analysis

To explore the segmented nucleoprotein of the novel Trematomous arenavirus, structures of the full coding regions (ORFs) of nucleoprotein segments were predicted using the Google Colab implementation of AlphaFold2 using MMseq2 with default settings (Mirdita et al., 2022). The resulting structural models were compared against the Protein Data Bank using the PDB search on the online Dali server (Holm, 2022) to identify which segments of arenavirus nucleoproteins the novel structures represented. Relevent structural superimpositions of novel nucleoproteins with those of a representative arenavirus nucleoprotein structure, *Lassa virus* (Protein Data Bank: 3MX5), and all figures of structures were prepared using UCSF ChimeraX (Pettersen et al., 2021).

### Testing effects of host factors on virome composition

The potential effects of host taxonomy and body mass group on virome composition were tested using various packages in R (v4.1.1). Standardised virome abundances were normalised and a distance matrix was created using the vdist function from the vegan package (Oksanen et al., 2022). Bray-Curtis dissimilarity as the distance measure and non-metric multidimensional scaling (NMDS) was performed on the distance matrix using the metaMDS function in vegan. Permutational multivariate analysis of variance (PERMANOVA) was used to test for statistical significance of the effect of host taxonomy (order and genus) and body mass group (high or low, see Table 1) on virome composition (presence and abundance) using the adonis2 function in vegan. Data were plotted using ggplot2 (Wickham, 2016).

### Virus nomenclature

Viruses were considered novel if they shared less than 90% RdRp or DdDp amino acid identity with their closest known relative, or less than 80% genome (nucleotide) identity with previously described virus species. This is a general classification based on ICTV guides for the classification of novel viruses for some taxonomic groups (Lefkowitz et al., 2018), as many have no or unclear genomic thresholds for proposing novelty, and the presumption that viruses with less than 90% amino acid identity in their most conserved proteins are unlikely to represent previously known species. Putative viruses were provisionally named according to either the common or scientific name of their host group.

### Data availability

Raw sequence reads are available on the NCBI Short Read Archive (SRA) under the BioProject accession PRJNA1088854. Virus genome sequences identified in this study are available under the GenBank accessions PP590684 – PP590779. Extended data outlining additional (non-polymerase) virus segments, lengths, and sequences from all of the viruses identified in this study, as well as R code and accompanying input data used to generate results and figures in this study can be found on GitHub: https://github.com/maybec49/Antarctic_Fish_Viromes.

## Results

### Virome diversity and abundance in Antarctic fish

Gill metatranscriptomes yielded approximately 61 – 119 million sequencing reads per library (Figure 1) and *de novo* assembly generated 473,258 – 782,113 contigs per library. Total virome reads represented 3.6×10^-5^ – 1.3% of the total sequencing reads (or 0.3 – 13,106 RPM) for each of the 22 RNA pools. *Antimora rostrata* (Gadiformes) with high body mass (> 2 kg) had the lowest viral abundance, while *Zoarcidae spp.* (Scorpaeniformes) in the low body mass group (< 0.11 kg) had the highest total virome abundance.

**Figure 1.**
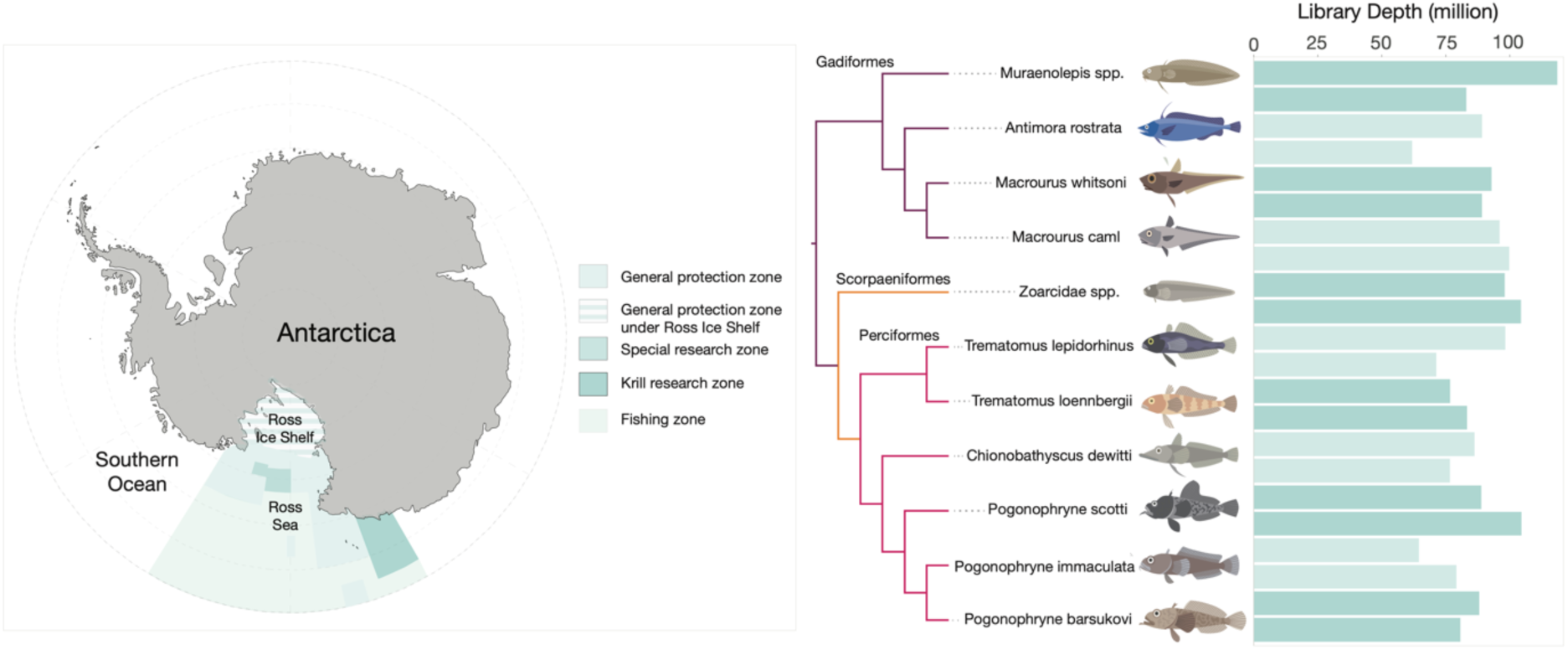
Ross Sea sampling site and phylogeny of fish hosts sampled for this study. Map of Antarctica with the Ross Sea region fish hosts that were sampled for this study highlighted in solid blue (left). Phylogeny of 11 fish species and family groups sampled for this study (middle). Branches are highlighted purple for Gadiformes, orange for Scorpaeniformes, and pink for Perciformes. Sequence library read depths for all 22 libraries (two per host group) in millions (right). Low body mass libraries are the first bars for each host, and high body mass is the second (underneath).

Antarctic fish viromes contained sequences from likely vertebrate-infecting viruses that could be assigned to one of 21 different viral orders or families (Figure 2). All 11 host species carried viruses from at least one taxonomic category. The number of virus taxonomic groups per host species ranged from one to eight, with an average of 4.4 per host. *Trematomus* (Perciformes) and *Muraenolepis* (Gadiformes) species carried the most (eight viral families) whereas the Perciformes species *C. dewitti* and *P. scotti* carried the least (one each). Only four of the 22 libraries contained no detectable fish virus sequences (low body mass *P. barsukovi* and *M. caml*, and high body mass *C. Dewitti* and *P. scotti*). Furthermore, transcripts from a total of 42 different viral polymerases or species were found across 15 RNA families, four DNA families, and one reverse-transcribed (RT) RNA family (Figure 2, Table 2). Perciformes carried 23 unique viruses, Gadiformes carried 15, and four were found in the Scorpaeniformes. Picornavirus sequences were the most commonly identified (in five host species), followed by astroviruses (in four host species). Also of note was the viruses identified were genetically distinct from any previously known viruses, representing entirely novel viromes. All of the viruses identified in this study shared 27 – 89% amino acid identity at most across all of the recovered virus segments (Table 2).

**Figure 2.**
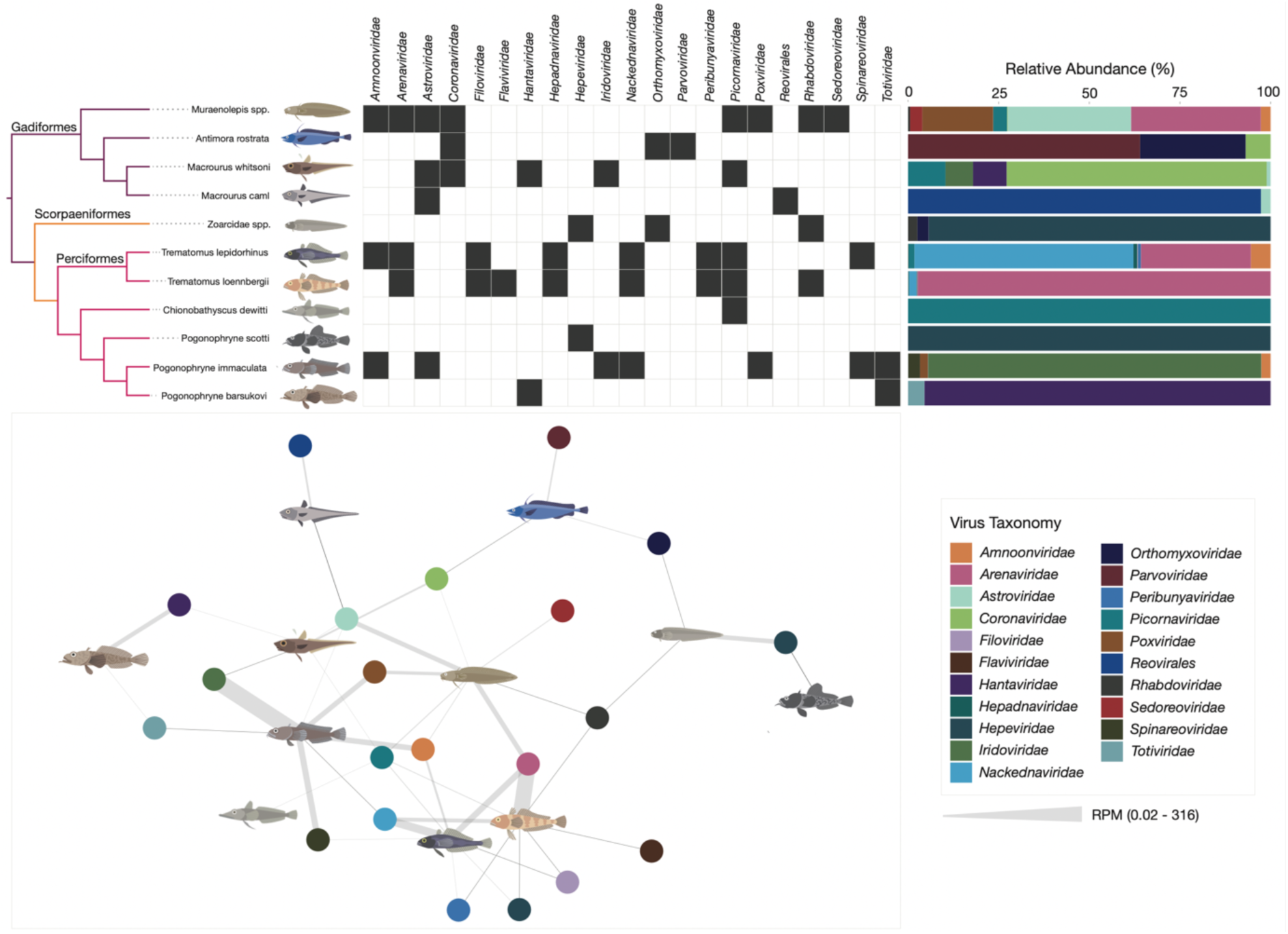
Antarctic fish viruses span 21 virus families and orders. Presence-absence map (top, middle) shows the presence of vertebrate viruses from 21 taxonomic groups in each host species group (top, left). Black boxes indicate the presence of the virus family/order in that host; white boxes indicate absence. Relative abundances (%) of each virus taxonomy in a host are shown (top, right). Network map showing connections between hosts and virus taxonomies (bottom, left). Fish hosts are shown by an illustration and virus families are indicated by coloured spheres. The width of grey connecting edges indicates abundance (in RPM).

**Table 2.**
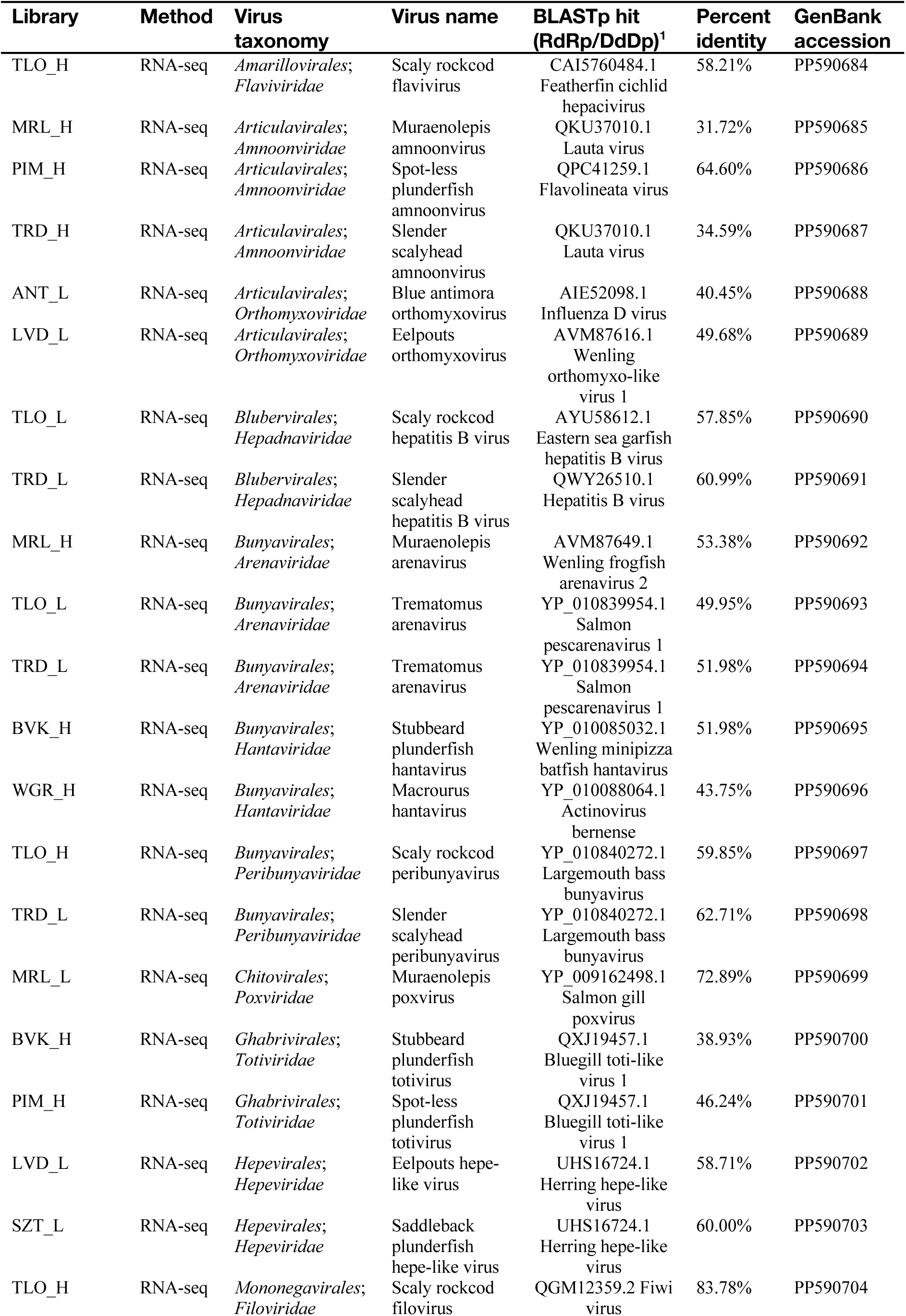
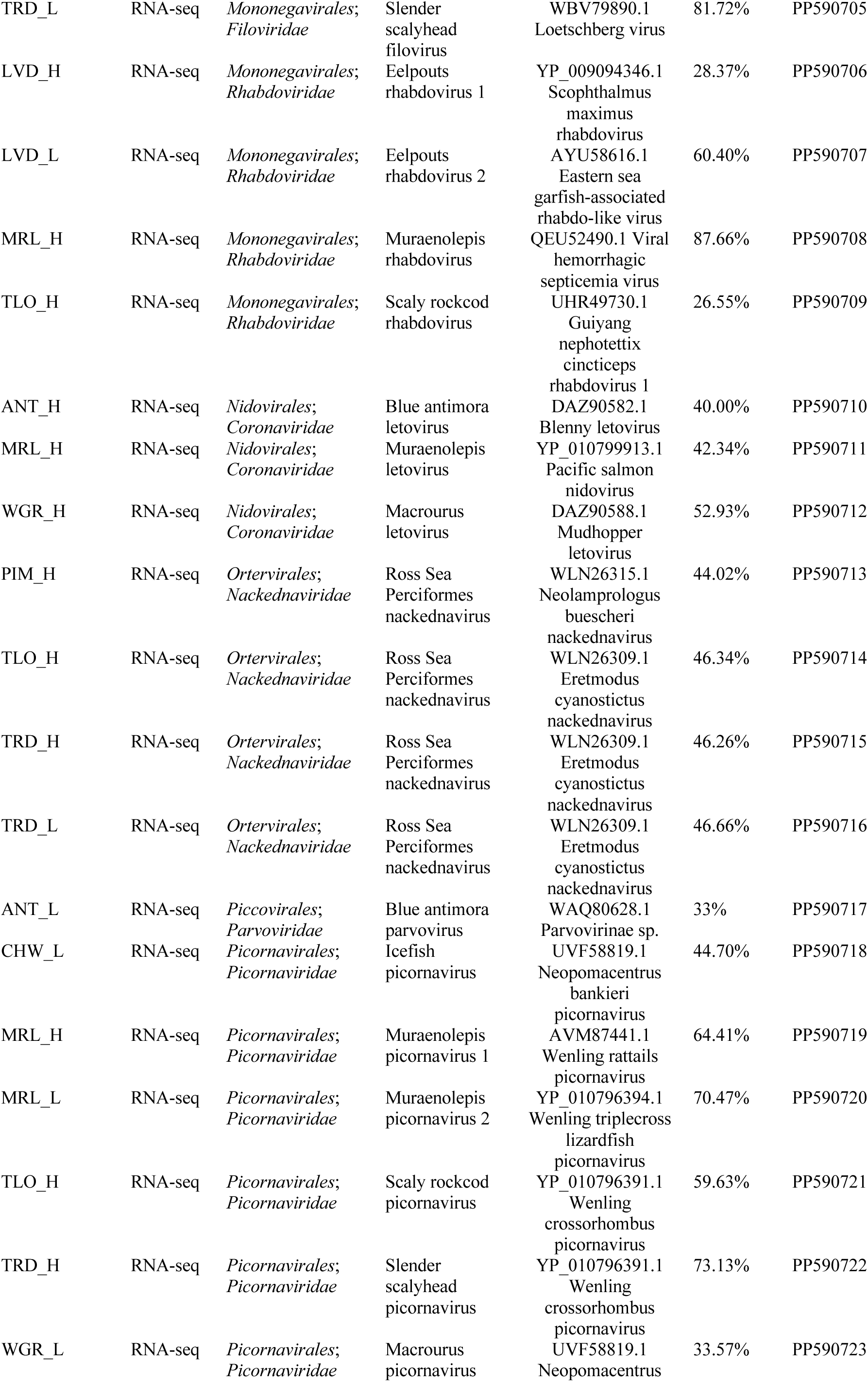
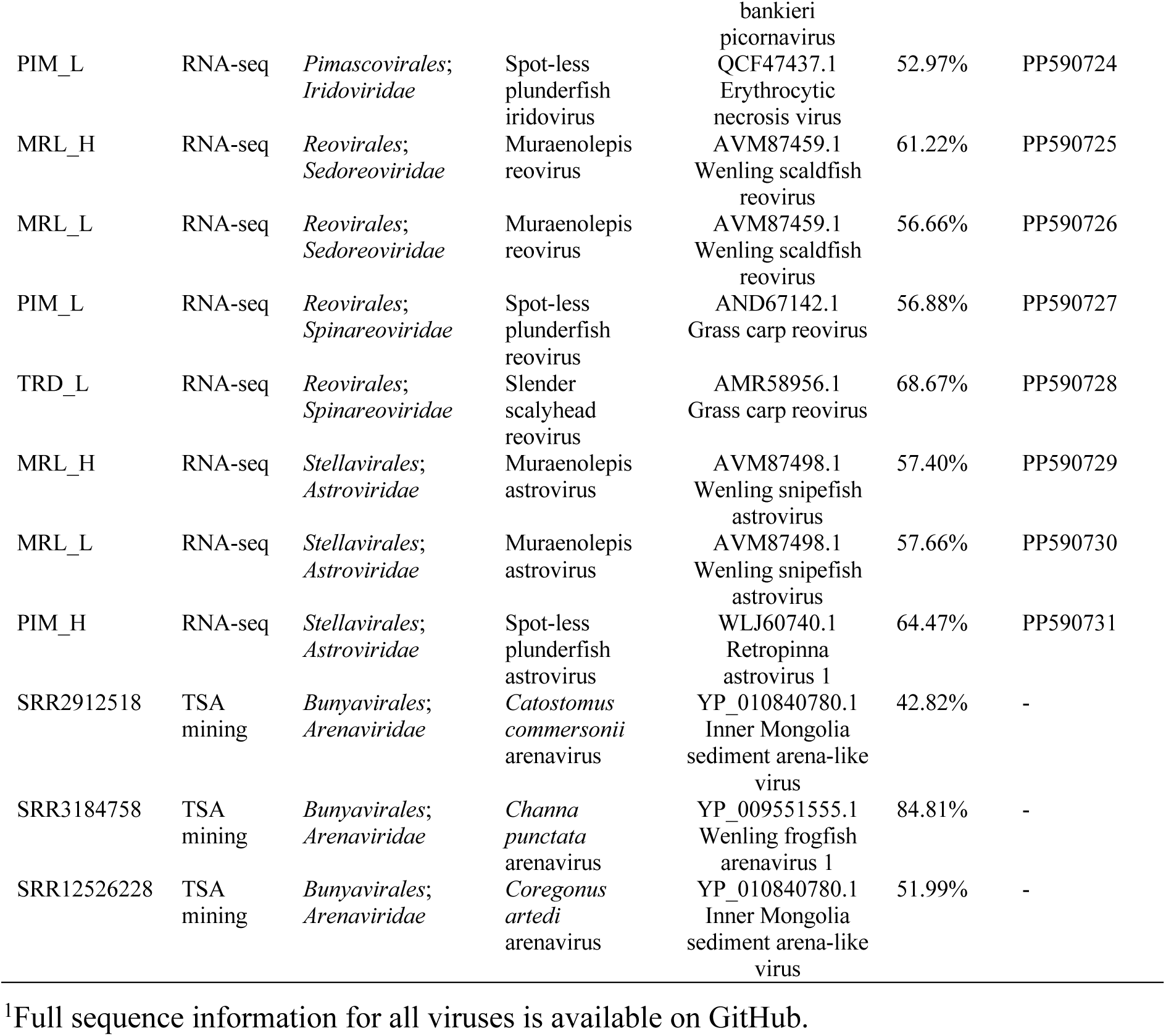
Putative virus polymerases identified in this study.

Vertebrate viral abundances were also variable across both host species and virus (Figure 2). Abundances ranged from 0.02 RPM to around 316 RPM. Perciformes and *Muraenolepis* spp. carried the highest number of viruses at the highest abundances. For example, viruses from the *Arenaviridae*, *Iridoviridae*, and *Nackednaviridae* were some of the most abundant in Perciformes (> 20 RPM, Figure 2). Other host species typically carried fewer than four different viruses and at lower abundances (< 5 RPM each). We also assessed the potential effects of host taxonomy and body mass grouping on virome composition. Neither body mass (p = 0.376) nor host order (p = 0.205) had a significant impact on virome composition, whereas host genus did (p = 0.013), although this could be confounded by the low sample sizes within each genus (Supplementary Figure 5).

Viral diversity in Antarctic fishes also closely mirrored that of fish hosts in other marine ecosystems. To compare this diversity across studies, we examined the number of virus taxonomic groups (representing virus families and orders) and unique viral sequences likely representing a species per host species alongside the total number of individuals sampled in recent virome studies of fishes (Supplementary Table 1 and Supplementary Figure 1 and 2). Generally, an increase in the number of individuals or host groups sampled corresponded with a greater number of unique viral species or sequences detected (R^2^ = 0.92, P < 0.01 for individuals; R^2^ = 0.85, P = 0.01 for host groups) (Supplementary Figure 1). However, we found no significant correlation between the number of individuals sampled per host and the number of virus taxonomic groups uncovered in this study (R^2^ = -0.06, P = 0.86). While direct comparisons between polar and tropical fish viromes could not be made due to fish viromes studies remaining limited, we could assess average diversity across studies. In the Ross Sea fishes, we identified viruses from 21 viral taxonomic groups, higher than the average of around 14 virus families/orders across recent studies, and all of which have previously been identified in fish (Supplementary Table 1 and Supplementary Figure 2). Antarctic fish harboured an average of 1.9 virus taxonomic groups per host, comparable to the average across studies of 1.8 (range: 0.33 to 2.2) and this observation was similar for the number of virus species per host (3.8) compared to the average (3.1; range: 1.2 to 5.5). Considering the influence of sample size on the number of viruses detected, we also examined the number of viruses per individual sampled. The average for Antarctic fishes was 0.17, consistent with the overall average in other studies (0.17; range: 0.14 to 0.27).

### Phylogenetic analysis of Antarctic fish RNA viromes

We inferred phylogenetic relationships of the novel RNA virus species using viral transcripts containing the highly conserved RdRp sequences. Overall, we identified 39 RdRps, representing 35 potentially unique species across 10 host species and all three host orders sampled (Figure 3 and 4).

**Figure 3.**
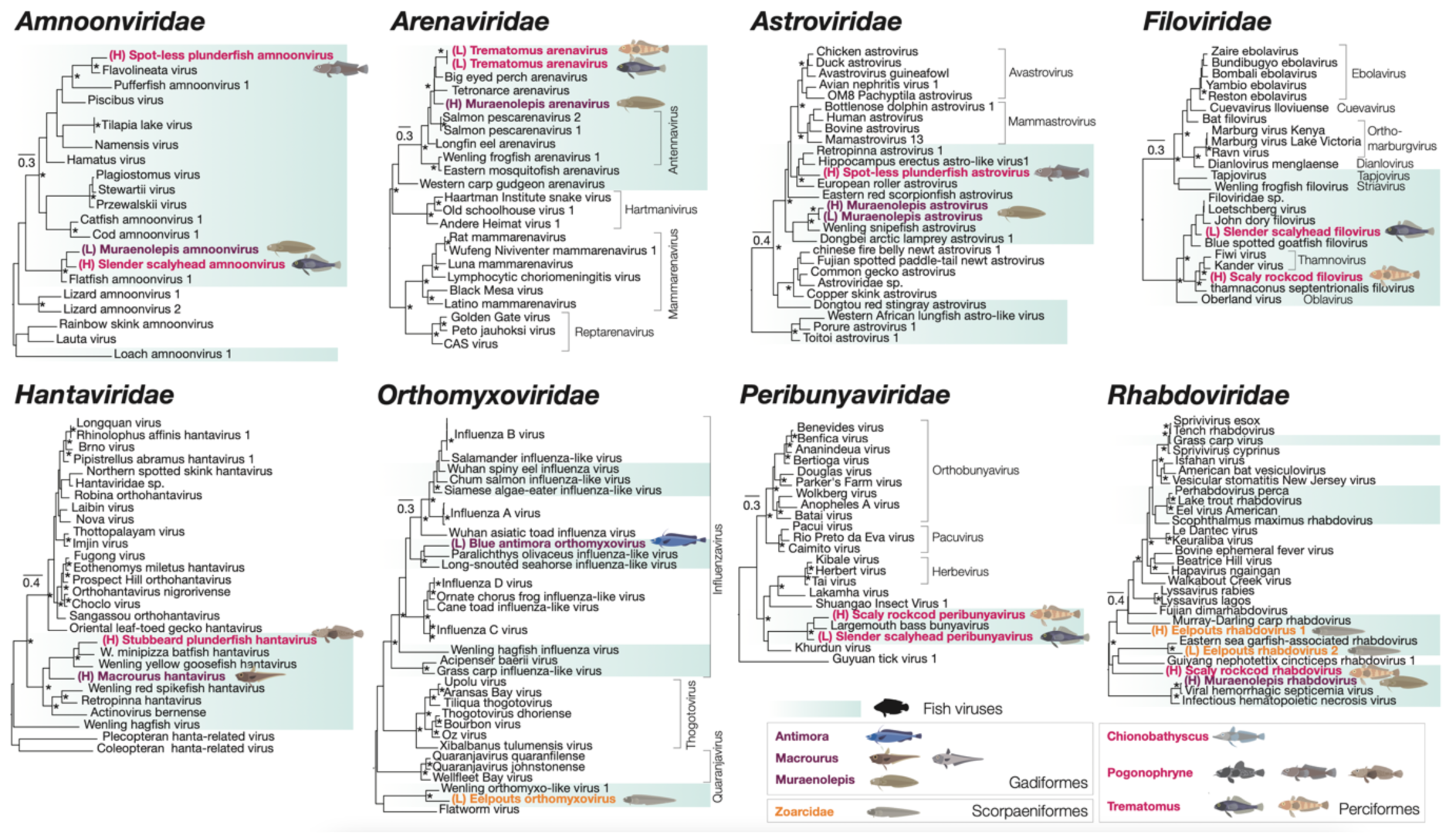
Phylogenetic analysis of negative-sense singe-stranded RNA viruses. Maximum likelihood trees of RdRp sequences of viruses from eight RNA virus families. Previously known and novel fish viruses are highlighted in blue. Novel viruses identified in this study are indicated by an illustration of their proposed host and highlighted based on the taxonomic order of their host: purple for Gadiformes, orange for Scorpaeniformes, and pink for Perciformes. Trees are rooted at midpoints and UF-bootstrap node support values ≥ 95% are denoted by an asterisk (*).

**Figure 4.**
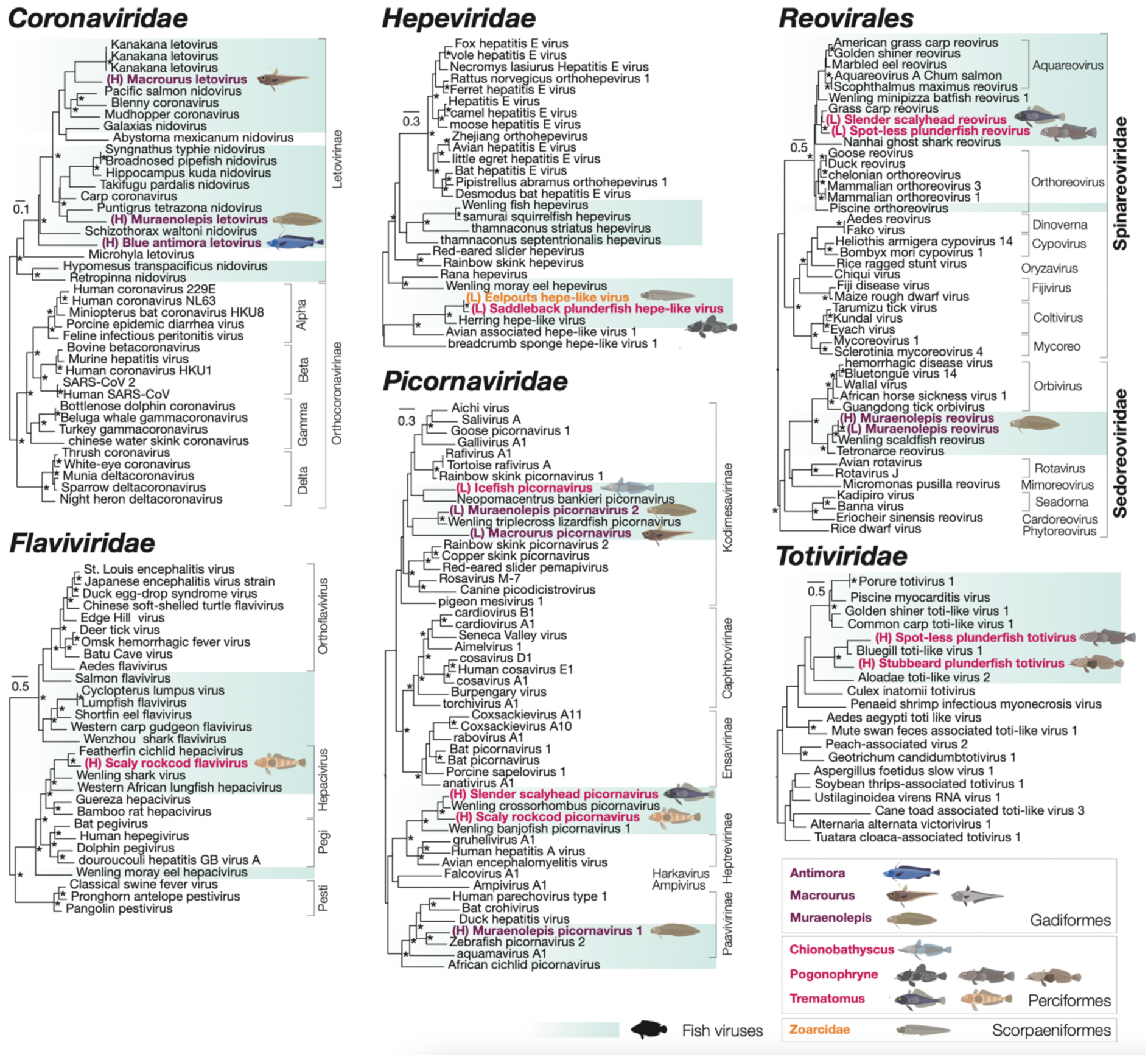
Phylogenetic analysis of positive-sense singe-stranded and double-stranded RNA viruses. Maximum likelihood trees of RdRp sequences of viruses from four +ssRNA and two dsRNA virus families. Previously known and novel fish viruses are highlighted in blue. Novel viruses identified in this study are indicated by an illustration of their proposed host and highlighted based on the taxonomic order of their host: purple for Gadiformes, orange for Scorpaeniformes, and pink for Perciformes. Trees are rooted at midpoints and UF-bootstrap node support values ≥ 95% are denoted by an asterisk (*).

The viruses spanned negative-sense single-stranded (ss)RNA families, including the *Amnoonviridae* (n=3 viruses), *Arenaviridae* (n=2), *Astroviridae* (n=3), *Filoviridae* (n=2), *Hantaviridae* (n=2), *Peribunyaviridae* (n=2), *Rhabdoviridae* (n=4), and *Orthomyxoviridae* (n=2) (Figure 3); positive-sense ssRNA families, including the *Coronaviridae* (n=3), *Flaviviridae* (n=1), *Hepeviridae* (n=2) and *Picornaviridae* (n=6); and the double-stranded (ds)RNA families: *Spinareoviridae* (n=2), *Spinareoviridae* (n=1), and *Totiviridae* (n=2) (Figure 4). All of these viruses fell into clades that contained previously known fish viruses (Figure 3 and 4, blue highlight). The *Picornaviridae* family had the largest expansion with six new species from five Perciformes and Gadiformes hosts being added. Notably, viruses from Perciformes were placed in 13 of the 15 families, excluding the *Coronaviridae* and *Sedoreoviridae* families.

### Negative single-stranded RNA viruses

Antarctic fish similarly harboured a number of -ssRNA viruses falling within the *Arenaviridae*, *Filoviridae*, and *Hantaviridae* that include viruses causing severe hemorrhagic diseases in a variety of hosts (Cobo, 2016) and in the *Orthomyxoviridae*, which includes respiratory disease-causing influenza and influenza-like viruses (Parry et al., 2020). Two new arenaviruses from Ross Sea fish, for example, provisionally termed Muraenolepis arenavirus and Trematomous arenavirus, shared ∼50% amino acid sequence identity with viruses such as *Salmon pescarenavirus 1* and *2*. Salmon arenaviruses have previously been associated with hemorrhagic symptoms in wild salmon (*Oncorhynchus tshawytscha*) (Mordecai et al., 2019). We also found rhabdoviruses in all three host orders studied here. One new virus, Muraenolepis rhabdovirus, shared over 87% amino acid identity with *Viral hemorrhagic septicemia virus*, which causes a potentially devastating hemorrhagic disease infecting a large number of species from the Perciformes and Gadiformes (Escobar et al., 2018). Other novel rhabdoviruses in scaly rockcod and eelpouts were more divergent in sequence, sharing less than 30% amino acid identity with other fish viruses linked to similar diseases, such as *Scophthalmus maximus rhabdovirus* (YP_009094346.1) (Zhang et al., 2007).

### Positive single-stranded RNA viruses

Viruses from the *Letovirinae* subfamily of the *Coronaviridae* were identified in a number of Gadiformes. The *Letovirinae* is comprised of viruses from aquatic vertebrates. Of note was the phylogenetic relationship between Macrourus letovirus, found here, to *Kanakana letovirus* from diseased New Zealand lamprey (*Geotria australis*) (Miller et al., 2021) (Figure 2). That these viruses formed sister-groups in the *Letovirinae* phylogeny is indicative of cross-order transmission over an uncertain time-scale. The diversity of picornaviruses was also expanded by viruses in both Gadiformes and Perciformes. Novel picornaviruses shared between 34 – 70% amino acid identity with viruses from other bony fish from China (Shi et al., 2018) and coral reef systems (Costa, Bellwood, et al., 2023), falling in previously established subfamilies such as *Kodimesavirinae* and *Paavivirinae*.

### Double-stranded RNA viruses

We identified dsRNA viruses from the order *Reovirales* and family *Totiviridae.* Totiviruses in two *Pogonophryne* hosts were both most closely related (sharing 39 – 46% amino acid identity) to Bluegill toti-like viruses found in deceased *Lepomis macrochirus* (bluegill) from Minnesota, USA (Sandlund et al., 2021). Interestingly, +ssRNA and dsRNA viruses from Perciformes and Gadiformes often fell into distinct clades or families. For example, both of the novel reoviruses from Perciformes hosts clustered in the *Spinareoviridae* family and were most closely related to viruses from the *Aquareovirus* genus, including Grass carp reovirus, which causes severe hemorrhagic disease in aquaculturally important species in China (Chen et al., 2013). Muraenolepis reovirus (from a Gadiformes species), on the other hand, fell into the *Sedoreoviridae* family most closely related to a likely benign virus, *Wenling scaldfish reovirus*, found in fish in China (Shi et al., 2018).

### Novel viruses DNA and reverse-transcribed RNA viruses

Two dsDNA viruses, one ssDNA virus, and a reverse-transcribed (RT) RNA virus, were found in Perciformes and Gadiformes, all falling into fish- or ectothermic vertebrate-associated groups of viruses (Figure 5, blue highlight).

**Figure 5.**
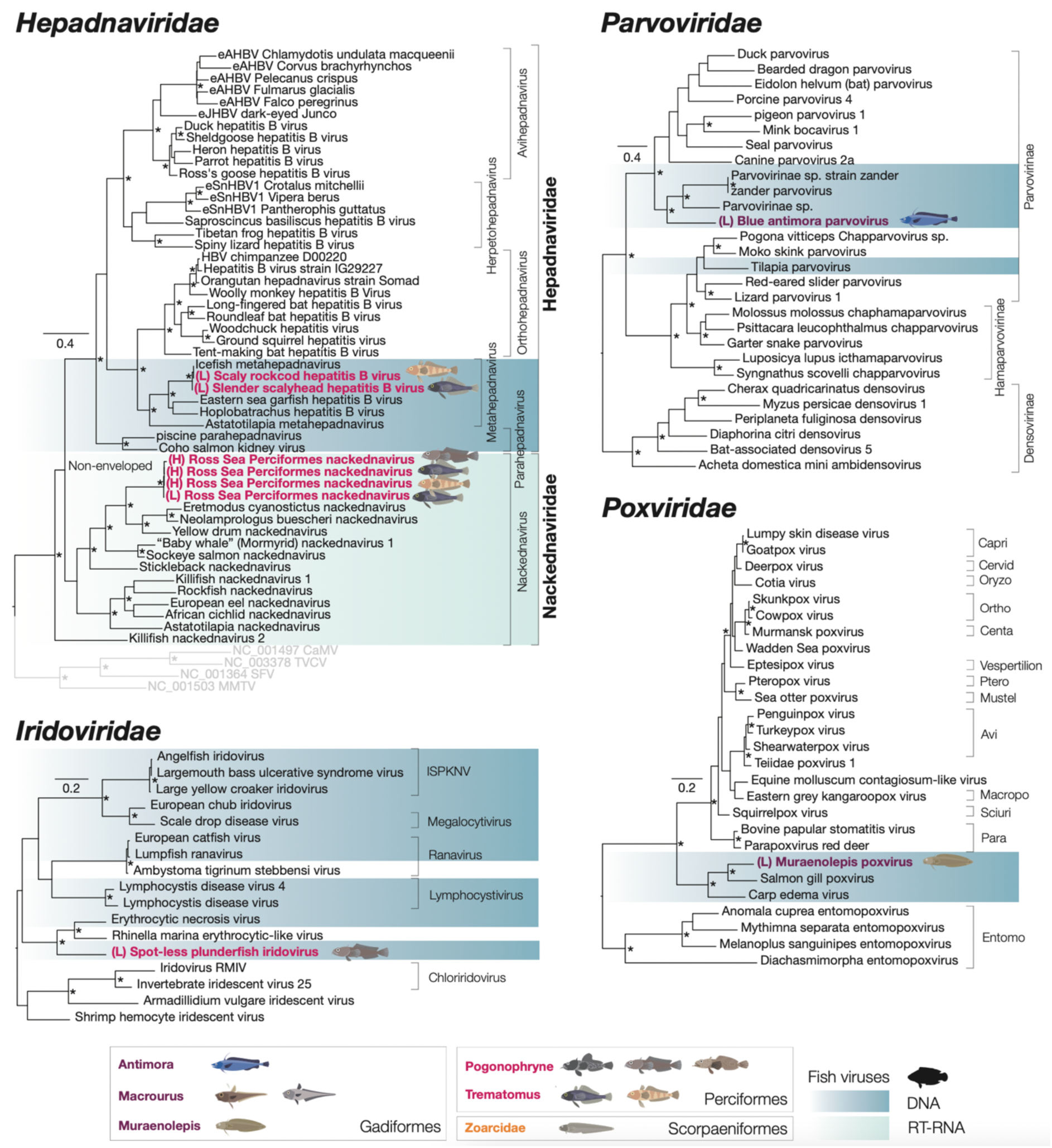
DNA and RT-RNA virus families expanded in this study. Maximum likelihood trees of DdDp or Pol proteins of viruses four dsDNA virus families and RdRps from a novel reverse-transcribed virus from the *Nackednaviridae*. The hepadnavirus phylogeny is based on a tree in (Lauber *et* al., 2017). Previously known and novel fish viruses are highlighted in blue. Novel viruses identified in this study are indicated by an illustration of their proposed host and highlighted based on the taxonomic order of their host: purple for Gadiformes, orange for Scorpaeniformes, and pink for Perciformes. Trees are rooted at midpoints and UF-bootstrap node support values ≥ 95% are denoted by an asterisk (*).

Hepadnaviruses (dsDNA) are responsible for hepatitis B infections in some mammals, although they have also been identified in fish (Dill et al., 2016). Nackednaviruses, on the other hand, are a more recently identified group of exogenous replicating viruses in fish that may have diverged from a common ancestor of hepatitis B viruses > 400 mya and have been hypothesised to help maintain the persistence of hepatitis B virus infections (Lauber et al., 2017). Two related hepadnavirus Pol genes falling in the *Metahepadnavirus* genus were found in both *Trematomus* species sharing 58-61% amino acid identity with the homologous sequences from hepatitis B viruses in fish (AYU58612.1) and amphibians (QWY26510.1) (Figure 5, Table 2). We also found a full 3,447 nt genome of a likely exogenous novel nackednavirus (RT-RNA) (see Supplementary Figure 1) tentatively named Ross Sea Perciformes nackednavirus (Figure 5). The nackednavirus was found in both *Trematomus* hosts as well as a *Pogonophryne* species (*P. immaculata*) and shared around 47% amino acid identity with *Eretmodus cyanostictus nackednavirus* (WLN26309.1) from a cichlid fish metagenome (Costa, Ronco, et al., 2023).

Two additional dsDNA viruses most closely related to disease-causing viruses of fish and other cold-blooded vertebrates were identified in the *Iridoviridae* and *Poxviridae* (Figure 5). Spot-less plunderfish iridovirus had ∼53% amino acid identity with *Erythrocytic necrosis virus* (QCF47437.1) which causes abnormalities of red blood cells and has been associated with mass mortalities in fish (Hershberger et al., 2009). Mauraenolepis poxvirus shared 73% amino acid identity with *Salmon gill poxvirus* (YP_009162498.1) which causes a potentially fatal respiratory disease in Atlantic salmon (Tartor et al., 2022).

Finally, we identified a ssDNA virus in the *Parvoviridae* in *Antimora rostrata* that exhibited ∼30% amino acid identity with Parvovirinae sp. (e.g. WAQ80628.1) from fish intestinal and metagenome samples (Reuter et al., 2022; Xi et al., 2023).

### Recent and historic cross-species transmission of viruses

Evidence of recent and likely historic host-jumping events was found between and within host orders, particularly in the Perciformes. We observed a probable instance of recent cross-species virus transmission within the *Arenaviridae*. Specifically, Trematomus arenavirus was identified in both *T. loennbergii* (scaly rockcod) and *T. lepidorhinus* (slender scalyhead) (Figure 3). The virus was 99% identical at the nucleotide level between the two hosts and did not fall below the abundance threshold which would be suggestive of index-hopping with abundances of 8.8 and 221.9 RPM. It also shared around 52% amino acid identity with the RdRp (L segment) of *Salmon pescarenavirus* (YP_010839954.1), representing a novel arenavirus most likely belonging to the *Antennavirus* genus of fish-infecting arenavirus (Pontremoli et al., 2019). The aforementioned Ross Sea Perciformes nackednavirus was also present in three different Perciformes and similarly appears to represent another more recent, and the most widespread, host-switching event. Examples of more historic host-jumping of viruses between Perciformes, particularly *Trematomus* species, were noted in the *Peribunyaviridae*, *Picornaviridae*, *Spinareoviridae*, *Totiviridae*, and *Hepadanaviridae* families due to the presence of closely related, but not identical viruses. Similar patterns of host-switching were not observed within the equally sampled Gadiformes order. Additionally, we did observed two likely examples of viruses jumping host orders, including that of amnoonviruses between *Muraenolepis* (Gadiformes) and slender scalyhead (Perciformes; *Trematomus*) and of hepeviruses between Eelpouts (Scorpaeniformes; *Zoarcidae*) and Saddleback plunderfish (Perciformes; *Pogonophryne*).

### Segmented nucleoprotein in *Trematomous* species expands genomic and structural diversity of the *Arenaviridae*

Of particular note was Trematomous arenavirus that possessed a novel genome and nucleoprotein structure (Figure 6 and Supplementary Figure 3). The virus shared more than 99% nucleotide identity between pools of two different species in the genus *Trematomous*: *Trematomus lepidorhinus* (Slender scalyhead) and *Trematomus loennbergii* (Scaly rockcod) at abundances of 8.8 and 221.9 RPM, respectively (Figure 6C). Trematomus arenavirus contained the typical three genome segments – with separate polymerase, glycoprotein, and nucleoprotein (NP) segments, found in other fish arenaviruses from the *Antennavirus* genus (Figure 6A) (Pontremoli et al., 2019). However, the nucleoprotein sequence in the novel virus was split over two ORFs as opposed to the canonical single ORF. The two ORFs appear to cover the nucleoprotein core domain in a 349 amino acid protein and the exonuclease domain in another 314 amino acid segment (Figure 6A and B). As this anomaly was present in the virus genome in both host species and at an abundance of > 0.1% in the library with a lower viral abundance (8.8 RPM), it suggests the split in the two nucleoprotein domains was not a result of a sequencing or assembly error nor a case of index-hopping.

**Figure 6.**
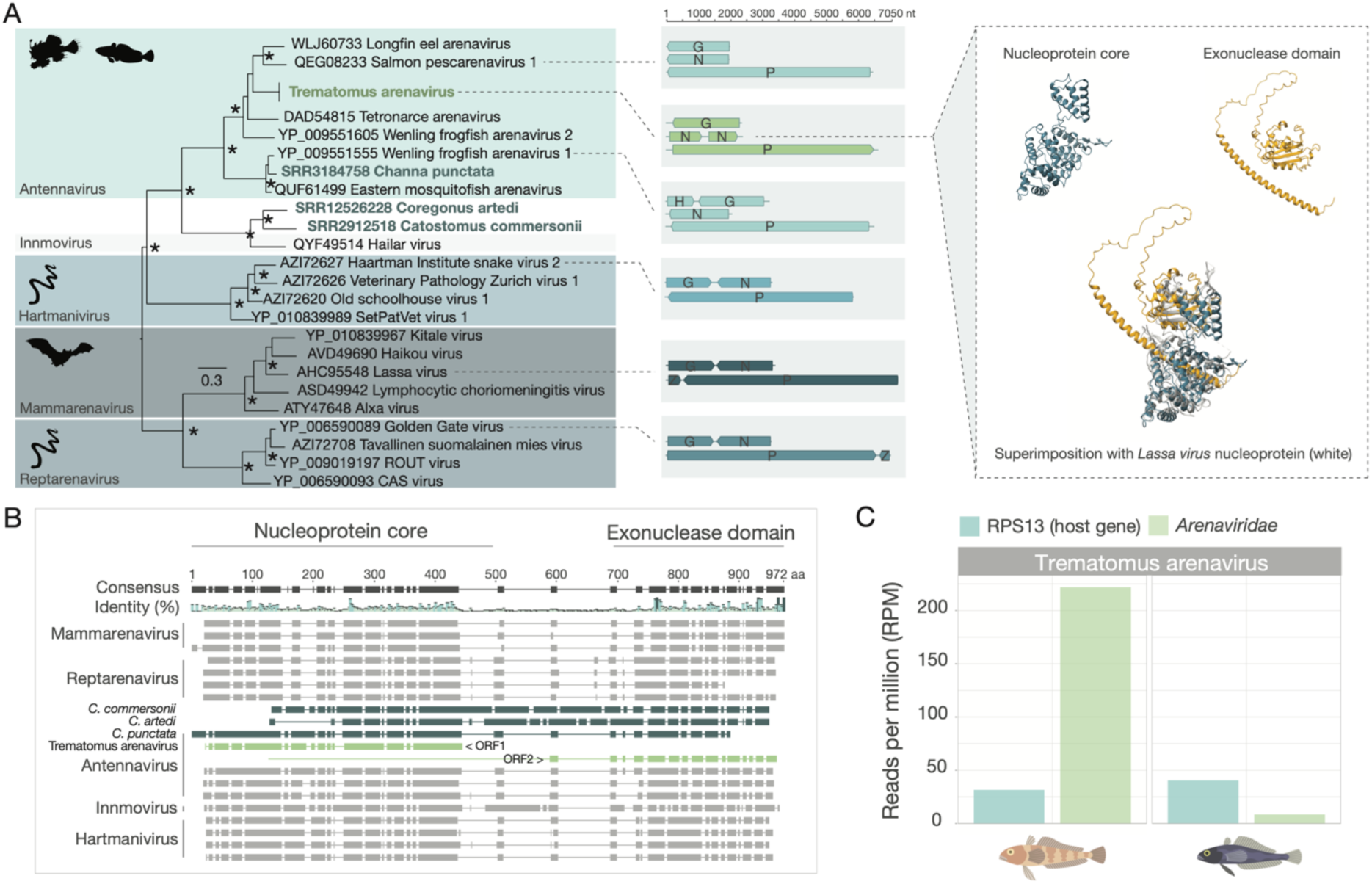
Trematomus arenavirus has a split nucleoprotein. A) Phylogeny (based on the RdRp sequence) of the *Arenaviridae* highlighted by genus (left). Trematomus arenavirus is highlighted in green and novel TSA-mined arenaviruses are highlighted in blue. Genome organisations of representative arenavirus genomes for each genus and their lengths (nt) showing the genomic structure diversity of the *Arenaviridae* (middle) and the protein structure of the segmented nucleoprotein core (blue) and exonuclease domain (orange) of Trematomus arenavirus, as well as a superimposition of these on the *Lassa virus* nucleoprotein structure (white) (Protein Data Bank: 3XM5), are shown (right). G = glycoprotein; N = nucleoprotein; P = polymerase (L segment); Z = Z protein; H = hypothetical protein. B) Protein alignment of nucleoproteins from arenavirus species. The linker between the nucleoprotein core and exonuclease domain is not well conserved across hosts. Note that the Trematomus arenavirus nucleoprotein (shown in green) does not form a contiguous ORF and instead consists of two ORFs encoding different segments of the full nucleoprotein (green). Trematomus arenavirus nucleoprotein segments are highlighted in green and novel arenaviruses nucleoproteins identified from the TSA by this study are highlighted in dark blue. C) Abundance (RPM) of host gene RPS13 (blue) compared to that of Trematomus arenavirus (green) in two *Trematomus* species.

At the structural level, the two nucleoprotein domains overlap closely with the nucleoprotein of *Lassa virus* (3MX5) (Figure 6A) with the exception of an approximately 60 amino acid α-helix structure in the N-terminus of the exonuclease domain with a high predicted local-distance difference test (Jumper et al., 2021) score (> 80) and which was not present in other nucleoprotein structures of known arenaviruses (Supplementary Figure 4).

To determine if there were other arenaviruses with similar unique genomic and structural variations, we employed an additional data mining approach to screen available transcriptome assemblies in the NCBI TSA database. Using polymerase, glycoprotein, and nucleoprotein sequences from Trematomous arenavirus as bait, we revealed an additional three fish-associated arenaviruses in *Channa punctata* (SRR3184758), *Coregonus artedi* (SRR12526228), and *Castomus commersonii* (SRR2912518) (Table 2, Figure 6A and B, and Supplementary Figure 4). Although partial segments could be recovered for all three segments for the three hosts, the viruses were not the closest relatives of Trematomous arenavirus (Figure 6A) nor were the nucleoprotein sequences similarly disrupted.

Partial nucleoproteins of the viruses from *C. comersonii* and *C. artedi* did reveal similar helices linking the nucleoprotein core and exonuclease domains (Figure 6B and Supplementary Figure 4A), providing evidence for this nucleoprotein structure as a unique feature of some aquatic arenaviruses.

## Discussion

Climate and geographical changes can drive the evolution and adaptation of hosts and their viruses (Danovaro et al., 2011). Fish from the Southern Ocean experienced a significant reduction in biodiversity as a result of continental shifts and cooling water temperatures (Clarke & Johnston, 1996; Eastman & Clarke, 1998), but the results of these events on virome diversity in these fish have not been explored. To our knowledge, studies on the viruses of these fish have remained limited to instances of single virus genera or groups (e.g. bacteriophages) in *Trematomus* species (Lopez et al., 2023; Van Doorslaer et al., 2018). By analysing gill metatranscriptomes of 11 groups of fish species collected from the Ross Sea region in Antarctica, we present entirely novel viromes that have greatly expanded the known aquatic vertebrate virosphere of Antarctica. In total, 42 unique fish viruses provide a new perspective on the virus diversity and evolution in these isolated, polar hosts.

Marine species richness significantly increases towards the equator (Rabosky et al., 2018). As the fishes of Antarctica are absent or rare in most other marine environments (Eastman, 2005), our primary goal was to investigate whether the reduction in fish biodiversity during past continental and climatic changes would be reflected in their viromes compared to those in more temperate or tropical environments. Ross Sea fish viromes harboured an assortment of RNA and DNA viruses spanning 20 families. Despite the apparent novelty of their viromes, all of their viruses fell within phylogenetic groups of viruses that have commonly been identified in other fish virome-wide studies (Costa et al., 2021; Filipa-Silva et al., 2020; Ford et al., 2024; Geoghegan et al., 2021; Perry et al., 2022). Recent studies have also begun characterising the viromes of fish in tropical ecosystems in more depth. The Great Barrier Reef in Australia, for example, boasts almost four times the number of fish species compared to the Southern Ocean (De’ath et al., 2012). Contrary to their limited host diversity, however, the observed viral family richness and number and types of viral families infecting Ross Sea fishes were comparable to those of this coral reef system (Costa, Bellwood, et al., 2023; Costa et al., 2024). In fact, similar questions have also been asked of penguins and their ticks from the Antarctic Peninsula, revealing RNA virome diversity akin to that in Australian waterbirds (Wille et al., 2020). Overall, these emerging results suggest that Antarctica’s isolated nature and low community richness have not adversely affected the viral diversity of its fauna.

Animal radiations, such as those observed in notothenioid fishes like the Perciformes sampled here, likely played a role in shaping viral diversity in Antarctica. As host species that have undergone rapid speciation would be more genetically and ecologically similar, many of the immunological, cellular, and environmental barriers normally limiting microbial host-switching would yet to be established, in turn increasing the rate of cross-species virus transmission (Charleston & Robertson, 2002; Longdon et al., 2014). As there is an inverse relationship between latitude and speciation in marine fish (Rabosky et al., 2018), it is possible that viral host-switching may be especially pronounced in marine environments with lower temperatures where endemism and speciation rates tend to be high. In the fish studied here, not only did the radiated Perciformes harbour more viruses than the equally sampled Gadiformes, but the presence of closely related viruses, including those from the *Arenaviridae*, *Nackednaviridae*, and *Spinareoviridae*, shared across multiple notothenioid hosts, and the general connectivity of their shared family-level virome network, highlights the occurrence of numerous recent or historical viral host-jumping incidents. A similar pattern is observed in the cichlid fish of Lake Tanganyika in Africa: the rapid radiation of these fish over the last 10 million years is associated with hosts that are very closely related and hence a high rate of cross-species virus transmission (Costa et al., 2024). This also illustrates that host evolutionary history, rather than environmental factors, is perhaps the most important determinant of rates of cross-species transmission and virus divergence.

In contrast to RNA viruses, on average DNA viruses tend to exhibit stronger congruency with their hosts’ evolutionary histories, indicative of long histories of co-divergence (Geoghegan et al., 2017; Peck & Lauring, 2018). Accordingly, the identification of closely related hepadnaviruses in two Perciformes strengthens the argument that reduced host barriers to virus transmission have enabled host-jumping that has led to a higher number of closely related viral species persisting despite the speciation of notothenioids occurring around 20 mya.

Continual global temperature changes are raising concerns about the imminent repercussions for biodiversity, species interactions, and infectious disease dynamics in aquatic species (Burge et al., 2014). Temperature is a key determinant in the distribution of ecothermic species and can impact interspecies interactions (Kordas et al., 2011) and disease susceptibility (Cohen et al., 2020; Páez et al., 2021). Rising water temperatures, for example, are forcing many aquatic species polewards (Danovaro et al., 2011; Nye et al., 2009). However, research addressing instances of viruses emerging as a direct consequence of altered host interactions in response to climate change remains limited.

Although animals can host a range of viruses asymptomatically (Wille & Holmes, 2020), the presence of viruses, like the novel iridovirus and poxvirus, closely related to known fish pathogens with high mortality rates (Jung et al., 2017; Nylund et al., 2008), should be of interest amid environmental shifts in regions with lower biodiversity, like the Southern Ocean (Duffy et al., 2016), where the risk of population declines due to emergent diseases is heightened by additional anthropogenic stressors such as overfishing and species introductions.

*Trematomus arenavirus*, found in scaly rockcod and slender scalyhead, also displayed a unique genomic variation. Unlike previously documented arenaviruses, *Trematomus arenavirus*, had its nucleoprotein sequence split over two non-overlapping ORFs. Arenavirus nucleoproteins (NPs) are well-conserved at both sequence and structural levels and are multifunctional, comprising a core domain that polymerises and protects the viral RNA and an exonuclease domain for degrading dsRNA, connected by “flexible” linkers (Papageorgiou et al., 2021). The split in the NP of *Trematomus arenavirus* occurred in the poorly conserved linker region between the two domains, which also contained an α-helix structure modelled with high confidence, uncommon in known arenaviruses and present in three of the four newly discovered fish arenaviruses. Gene fusion and fission – the joining or splitting of multiple genes/protein domains (Marsh & Teichmann, 2010) – are fundamental evolutionary processes observed in cellular organisms (Snel et al., 2000). However, these processes have not been well studied in viruses. The mechanisms behind this domain split and the well-structured helix, potentially illustrating adaptation to extreme temperatures affecting protein folding (Feller, 2018), remain unknown, as does any interaction between the split domains.

Our study has a number of limitations. Pooling samples into groups assists in the efficiency and likelihood of virus discovery as often a large number of individuals can be screened with low viral yield (Y. Q. Li et al., 2023), but this also limits conclusions that can be drawn due to often low resulting library (sample) numbers. For instance, the capacity to estimate the prevalence of viruses in a host population or quantify the frequency of various evolutionary events is impacted. Tissue tropism is also frequently exhibited by viruses (Watashi & Wakita, 2015), and thus the choice of tissue examined will necessarily influence what viruses can be uncovered. These viromes represent that of a single tissue – the gills. Nevertheless, gills are a common site of virus entry and infection in fish (Kim & Leong, 1999) and analysis of viromes using multiple tissue types has shown that 94% of the viruses could be found in the gills alone (Costa, Ronco, et al., 2023), so it is likely that a majority of the viruses infecting these Antarctic fish could be successfully identified here using gills.

In sum, we explored the implications of isolation and host adaptation in extreme environments on virus evolution by analysing the gill-associated viromes of fish from the Ross Sea, Antarctica. Our findings revealed an abundance of RNA and DNA viruses, greatly expanding the known virosphere of these polar species. Overall, their virome diversity mirrored that of fish from more diverse and warmer marine environments, suggestive of the maintenance of high viral diversity in these taxonomically restricted populations. While we provide a more extensive catalogue of the known viruses of Antarctic fishes, future research would benefit from broadening the scope to better understand the effects of host evolution and adaptation to polar environments on the evolution of their viruses or explore more directly whether viral diversity or virome connectivity differs greatly at the poles compared to temperate or tropical environments.

## Acknowledgements

We acknowledge the crew of the San Aotea II, the NZ/MPI observer Nigel Hollands, the CCAMLR international observer (CapFish) Chuma Sijaji, Mike Prasad (the scientist onboard), Jennifer Devine (the scientist on shore), and funding provided by Fisheries New Zealand under MPI project ANT2019-01C. We also thank Hamish Thompson for his Antarctic fish species illustrations.

