## Supplementary figures and images for "Viromes of Antarctic fish resembles the diversity found at lower latitudes"

### Supplementary Figure 1

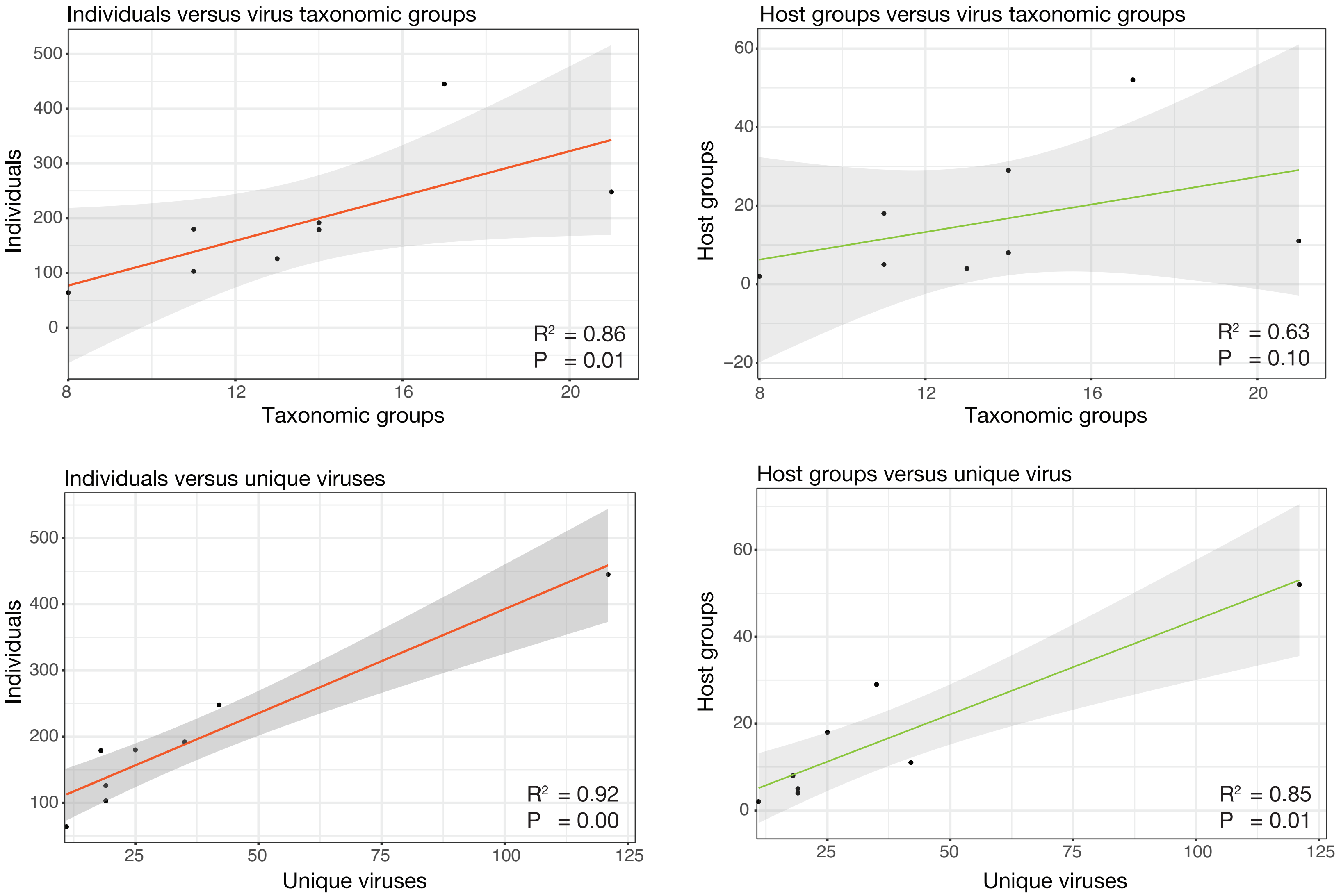

### Supplementary Figure 2

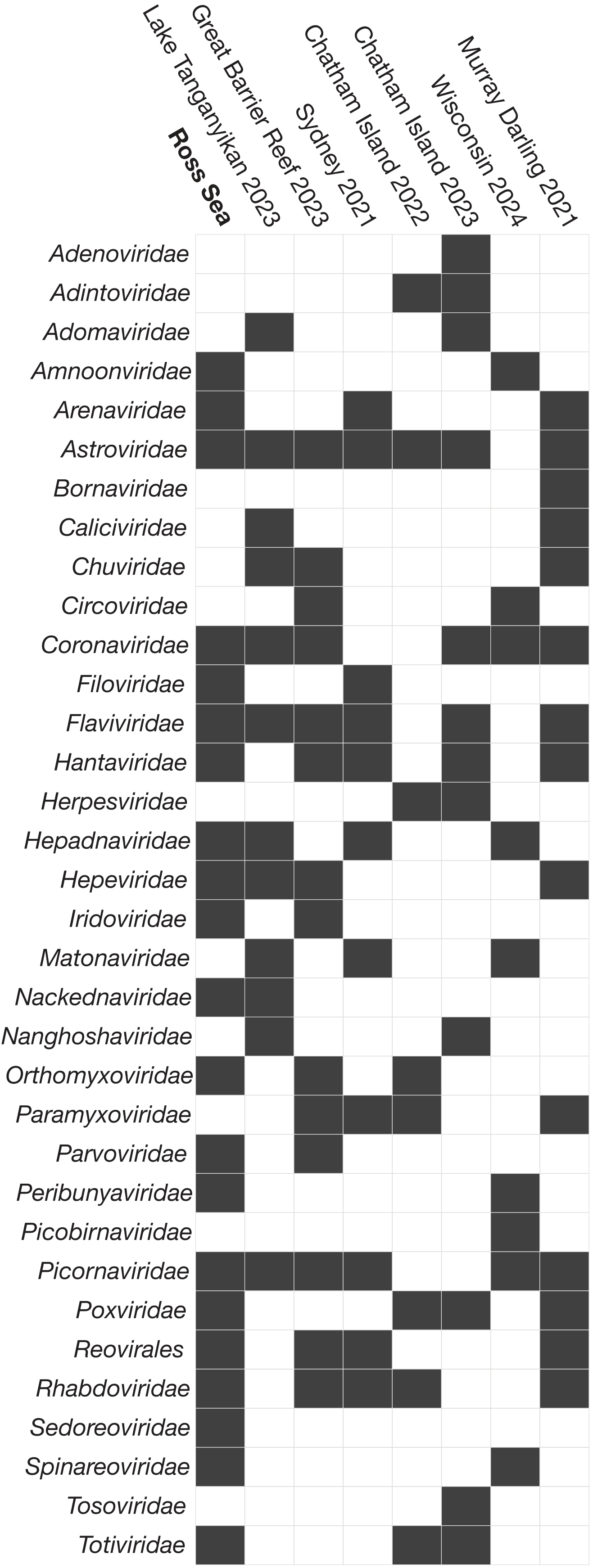

### Supplementary Figure 3

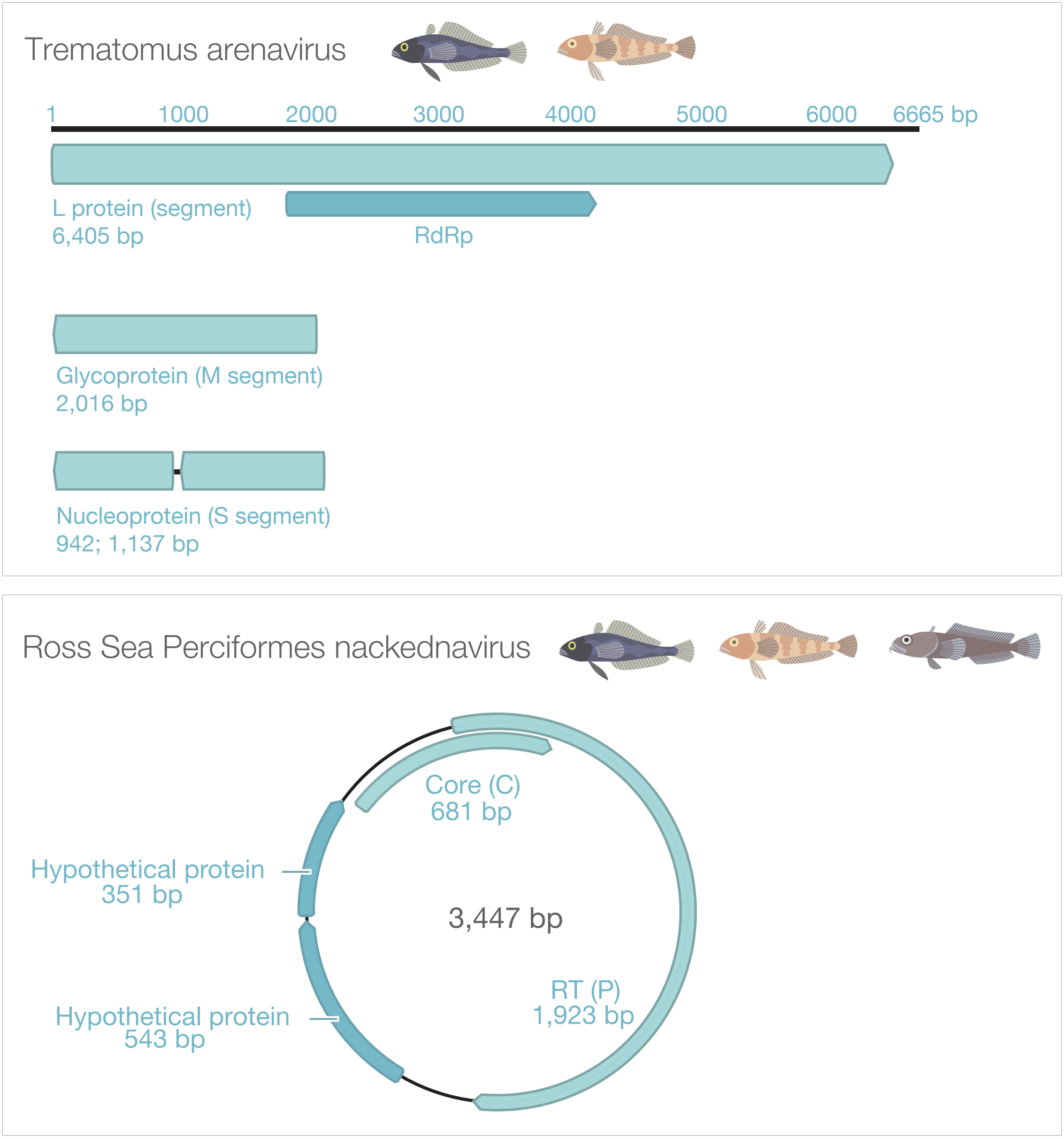

### Supplementary Figure 4

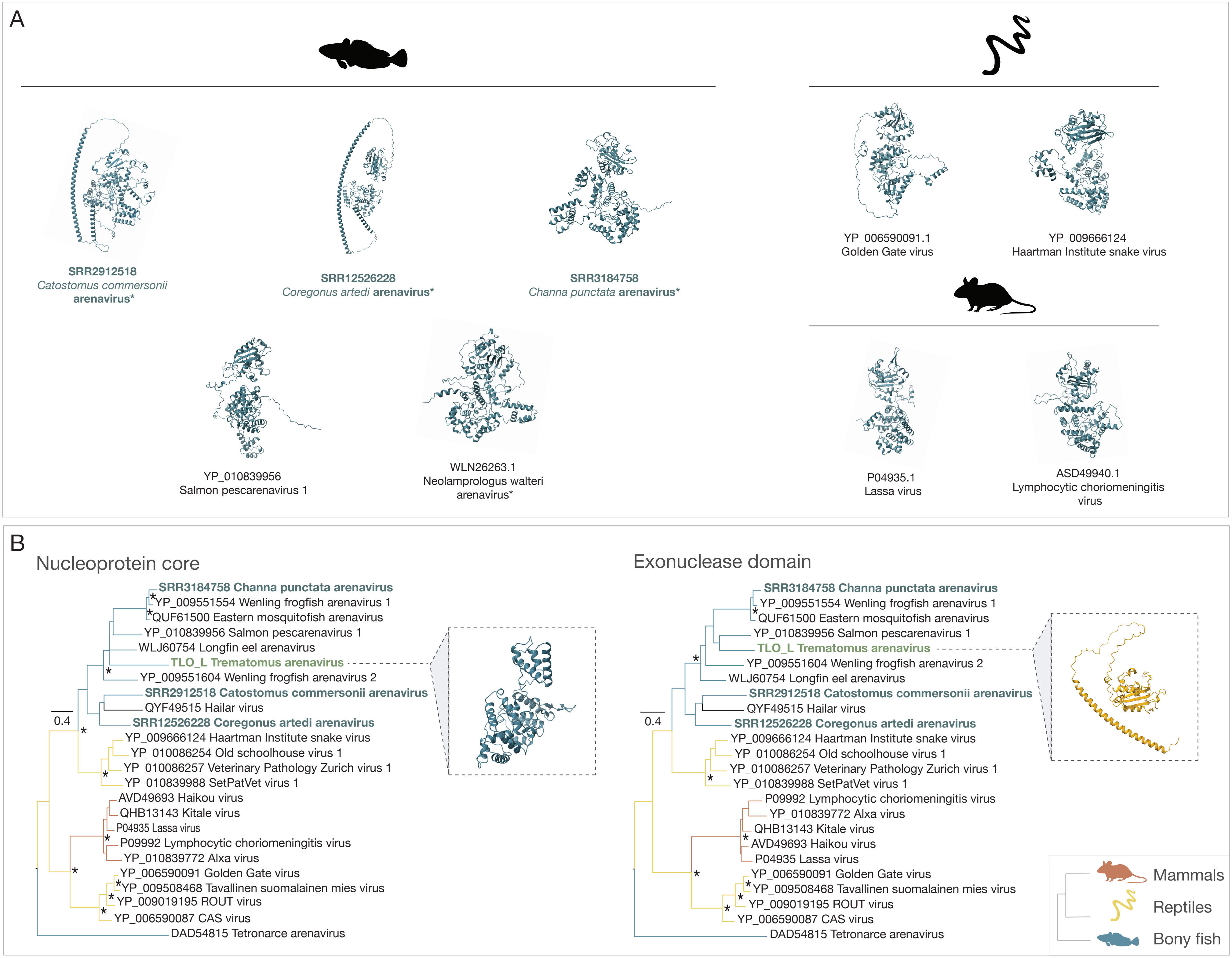

### Supplementary Figure 5

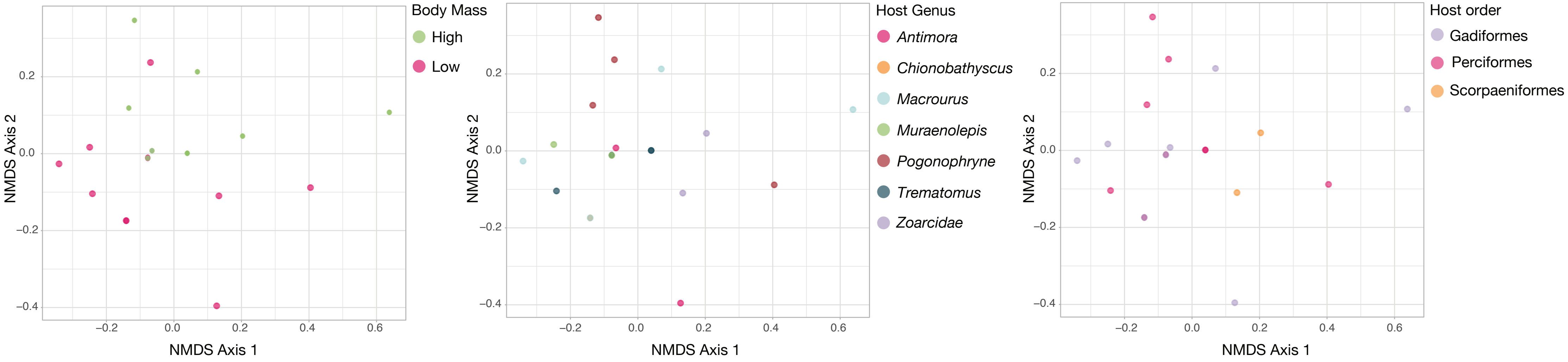
