## Supplementary Table 1 for "Viromes of Antarctic fish resembles the diversity found at lower latitudes"

**Supplementary Table 1.** Virus discovery in fish viromes^1^.

| **Location of study** | **Number of individuals^2^** | **Hosts with vertebrate viruses** | **Virus taxonomic groups** | **Unique viruses** | **Viral groups/hosts** | **Viral species/hosts** | **Viral groups/individuals** | **Viral species/individuals** | **Paper** |
| --- | --- | --- | --- | --- | --- | --- | --- | --- | --- |
| Ross Sea, Antarctic | 248 | 11 | 21 | 42 | 1.9 | 3.8 | 0.08 | 0.17 | This study |
| *Perciformes* | *118* | *6* | *16* | *22* | *2.7* | *3.7* | *0.14* | *0.19* | *This study* |
| *Gadiformes* | *121* | *4* | *11* | *14* | *2.8* | *3.5* | *0.09* | *0.12* | *This study* |
| Lake Tanganyikan, Africa | 445 | 52 | 17 | 121 | 0.33 | 2.3 | 0.04 | 0.27 | Costa et al., 2023 |
| Great Barrier Reef, Australia | 192 | 29 | 14 | 35 | 0.48 | 1.2 | 0.07 | 0.18 | Costa et al., 2023 |
| Murray-Darling Basin, Australia | 179 | 8 | 14 | 18 | 1.8 | 2.3 | 0.08 | 0.1 | Costa et al., 2021 |
| Sydney, Australia | 180 | 18 | 11 | 25 | 0.61 | 1.4 | 0.06 | 0.14 | Geoghegan et al., 2021 |
| Chatham Island, New Zealand | 126 | 4 | 13 | 19 | 3.3 | 4.8 | 0.10 | 0.15 | Grimwood et al., 2023 |
| Chatham Island, New Zealand | 64 | 2 | 8 | 11 | 4 | 5.5 | 0.13 | 0.17 | Perry et al., 2022 |
| Wisconsin, USA | 103 | 5 | 11 | 19 | 2.2 | 3.8 | 0.11 | 0.18 | Ford et al., 2024 |

^1^In studies published from 2020 to 2024 reporting numbers of individuals and host species collected.

^2^Total number of individual samples collected, including those with no vertebrate viruses.
